# Human Brain Endothelial Cell Seeded on Inner Surface of Alginate Hollow Microfibers

**DOI:** 10.1101/2023.01.04.522758

**Authors:** Saurabh S. Aykar, Nima Alimoradi, Isaac S. Petersen, Reza Montazami, Amanda L. Brockman, Nicole N. Hashemi

## Abstract

Barrier functionality of the blood-brain barrier (BBB) is provided by the tight junctions formed by a monolayer of the human brain endothelial cells (HBECs) internally around the blood capillaries. To mimic such barrier functionality in vitro, replicating the hollow tubular structure of the BBB along with the HBECs monolayer on its inner surface is crucial. Here, we developed an invasive microfluidic technique to obtain the HBECs monolayer on the inner surface of alginate-based hollow microfibers. The HBECs were seeded on the inner surface of these microfibers using a custom-built microfluidic device. The seeded HBECs were monitored for 9 days after manufacturing and cultured to form a monolayer on the inner surface of the alginate hollow microfibers in the maintenance media. A higher cell seeding density of 217 cells/mm length of the hollow microfiber was obtained using our microfluidic technique. Moreover, high accuracy of around 96 % was obtained in seeding cells on the inner surface of alginate hollow microfibers. The microfluidic method illustrated in this study could be extrapolated to obtain a monolayer of different cell types on the inner surface of alginate hollow microfibers with cell-compatible ECM matrix proteins. Furthermore, it will enable us to mimic a range of microvascular systems in vitro by closely replicating the structural attributes of the native structure.

## Introduction

Blood-brain barrier (BBB) comprises majorly of endothelial cells that line the interior of blood capillaries to form tight junctions which provide barrier functionality. ^1, 2^ The pericyte and astrocyte cells lay on the exterior of the endothelial cell layer and collectively maintain the selective permeability and integrity of the BBB. ^3, 4^ For the past decade, there has been extensive research to mimic the barrier function of the BBB using various high throughput microfluidic devices for several applications such as understanding the pharmacokinetics of different drugs. ^5–10^ The state-of-the-art devices replicate the BBB using a 2D surface where the endothelial cells are cultured on a porous membrane or on a static 3D scaffold that fails to impart the change of volume phenomena such as inflammation in the barrier model. ^11–18^ Although these devices enable us to investigate the selective permeability of the BBB to a certain length, they all lack the inclusion of physiological and physiochemical effects of dynamic 3D microstructure on the development and homeostasis of the BBB.

The BBB has a hollow tubular structure with endothelial cells lining the interior surface of the blood capillaries providing the barrier functionality. ^19–21^ The barrier functionality highly depends on the structure of the BBB, which swells or contracts to modulate the permeability of the barrier. To closely bio-mimic the essential attributes of BBB in vitro, it is crucial to replicate the native structure along with the monolayer formation of endothelial cells on its internal surface. The hydrogel-based 3D hollow scaffold will enable the replication of the in vivo like structure by additionally providing properties such as controlled swelling and permeation. ^22–24^ We have previously developed a microfluidic device that manufactures alginate hollow microfibers with cells invasively encapsulated or seeded in the microfiber walls or on the internal surface, respectively. ^25–27^ One of the novelties of this technique is the simultaneous seeding of cells during the manufacturing of hollow microfibers. Moreover, since the scaffold cannot be coated with the extracellular matrix prior to cell seeding as both occur simultaneously, we have also developed a technique to uniquely grow the human brain endothelial cells (HBEC) seeded on the internal surface of the alginate hollow microfibers. To this date, we know of no reports that have demonstrated the establishment of an HBEC monolayer on the inner surface of a 3D hollow dynamic scaffold to mimic the endothelium of the BBB.

In this study, we expound on the procedure of obtaining a monolayer of HBECs on the inner layer of alginate hollow microfibers over a prolonged period (9 days) in the culture to explicitly mimic the endothelial cell barrier. It is known that some cell type exhibits better adherence and viability to a specific extracellular matrix (ECM) proteins. ^28^ HBECs adhere well to entactin-laminin collagen IV and gelatin-coated flat surfaces leading to higher cell viability as compared to other ECM proteins. ^29, 30^ This study translates the HBEC adherence properties from a flat surface to a 3D robust surface of the alginate hollow microfibers simultaneously during their manufacturing while maintaining excellent cell viability. Subsequently, the future studies aim to conduct experiments to analyze the adherence and tight junction proteins along with some essential transporter proteins expressed by the HBEC monolayer obtained by this technique. This technique is not confined to obtaining HBEC monolayer and can be used as a platform to seed and grow different cell types on the inner surface of alginate hollow microfibers, which could be used to mimic different microvascular system in the body for drug delivery applications.

## Materials and methods

### Cell culture

Human brain endothelial cells (HBEC) (CRL-3245, ATCC, Manassas, VA) were used for cell seeding on the inner surface of alginate hollow microfibers simultaneously during their manufacturing. HBEC were cultured according to the company protocol in a maintenance media (MM) comprised of the following components: 90 % DMEM-F12 medium (30-2006, ATCC), 10 % Fetal Bovine Serum (FBS) (30-2020, ATCC), and 40 μg ml.^−1^ endothelial cell growth supplement (ECGS) (356006, Corning, NY). T-75 flasks (Fisher Scientific) coated with 0.1 % gelatin (PCS-999-027, ATCC) for at least 2 h were used for culturing HBEC. The culture was stored in the incubator and maintained at 37 °*C* and 5 % CO_2_ until 90% confluency was reached. The cell density used for seeding was 10 × 10^6^ *cells mL*^−1^.

### Cell seeding technique

The detailed design and manufacturing of the microfluidic device used for seeding cells on the inner surface of the alginate hollow microfibers are reported in our previous articles.^25, 26^ Briefly, a microfluidic design consisting of five inlet channels converging on a central channel and two sets of chevrons were micro-milled on polymethyl methacrylate (PMMA) chips and subsequently bonded together using thermally-solvent assisted bonding technique to obtain a microfluidic device. 2 w/v % Sodium Alginate (Sigma-Aldrich) in 0.1% gelatin water (PCS-999-027, ATCC) was mixed with 10 μg mL^−1^ entactin-laminin collagen IV (ECL) (08-110, Sigma-Aldrich) and the resulting solution was infused through the cladding inlets as a precursor. 10 w/v % polyethylene glycol (PEG) (8-18897, Sigma-Aldrich) in 0.1% gelatin water was infused through the sheath inlets as a template fluid. 20 w/v % of PEG was mixed with the HBEC suspended in the MM in an equal ratio to obtain a final concentration of 10 w/v % of PEG and was infused through the core channel. All solutions were prepared 12 h before experimentation and stored in the incubator with constant parameters of 37 °*C* and 5 % CO_2_. The solutions were pumped with a flow rate ratio (FRR) of 200: 250: 150 (core: cladding: sheath) *μL min*^−1^ using syringe pumps (Cole-Parmer, IL, USA). 10 w/v % Calcium Chloride Dihydrate (C79-3, Thermo Fisher Scientific) in DI water was used as a polymerizing agent as well as a collection bath. The HBEC seeded alginate hollow microfibers were collected and stored individually in separate wells in a 6-well plate containing MM. They were stored in the incubator for nine days after manufacturing, and the MM media was changed every three days.

### Imaging and analysis

The HBEC seeded alginate hollow microfibers were imaged under the confocal microscope (Zeiss) on different days after manufacturing, day 0, day 1, day 3, day 6, and day 9. The images were captured using a 10x and a 20x objective to ascertain the formation of the HBEC monolayer on the inner surface of the alginate hollow microfibers for 9 days after manufacturing. The seeded cells, on day 0, were counted using ImageJ software. The cells seeded per unit length in the hollow region and in the walls of the microfibers were plotted for comparison.

## Results and discussion

Human Brain Endothelial cells (HBEC) were invasively seeded on the inner surface of the alginate-based hollow microfibers using a previously built microfluidic device. The schematic illustration of the microfluidic cell seeding process is shown in **Figure 1**. The cells were monitored for 9 days after seeding to investigate cell adherence and growth on the inner surface of these alginate hollow microfibers. **Figure 2** shows the images of the HBECs seeded alginate microfibers on day 0 and on day 1, which are immediately and 24 h after manufacturing, respectively. The seeded cells, both in the hollow region and in the walls, were counted using ImageJ and were plotted per unit length of the microfiber, as shown in **Figure 3**. The plot depicts that a higher cell seeding density per unit length of 217 cells/mm was obtained. The higher cell seeding density is critical for the initiation of the growth of the cells within the hollow microfibers. This early growth initiation of the cells due to higher initial density was also observed from the day 1 images. The number of cells/mm that was leaked into the walls of the hollow microfibers during manufacturing was measured to be 8 cells/mm. This means around 96 % accuracy was obtained in seeding cells on the inner surface of the alginate hollow microfibers. This indicates that very few cells were encapsulated in the walls of the hollow microfibers in comparison to the ones seeded in the hollow region, and a significant difference was observed (p < 0.05).

**Figure 1.**
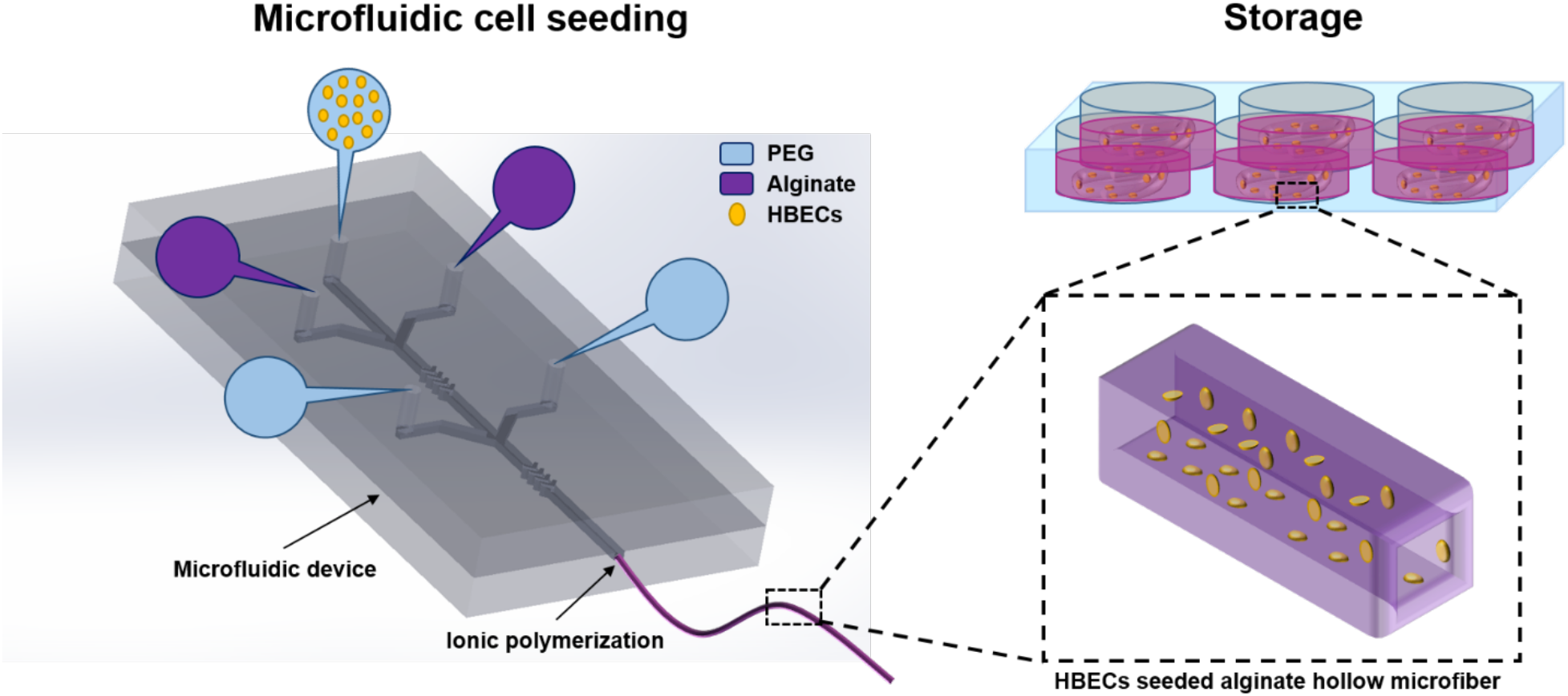
Schematic representation of the microfluidic approach to seed and obtain human brain endothelial cell (HBEC) monolayer on the inner surface of the alginate hollow microfibers.

**Figure 2.**
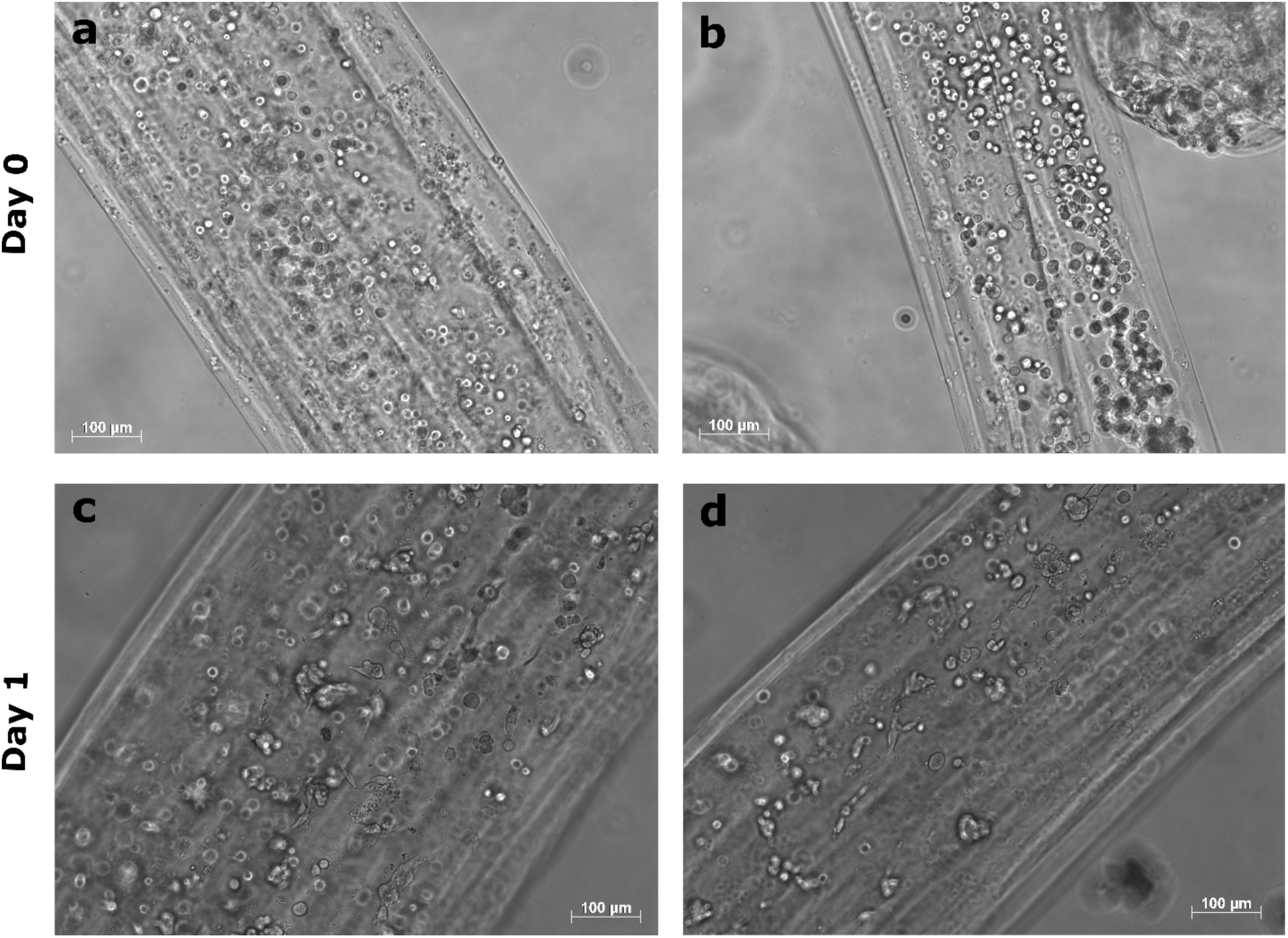
Microscopic brightfield images of the HBEC seeded alginate hollow microfibers captured after microfluidic manufacturing. **a** & **b** show the HBECs seeded immediately after manufacturing (Day 0). **c** & **d** depict the initiation of HBEC processes on the inner surface 24 h (Day 1) after manufacturing. All scale bars are 100 μm.

**Figure 3.**
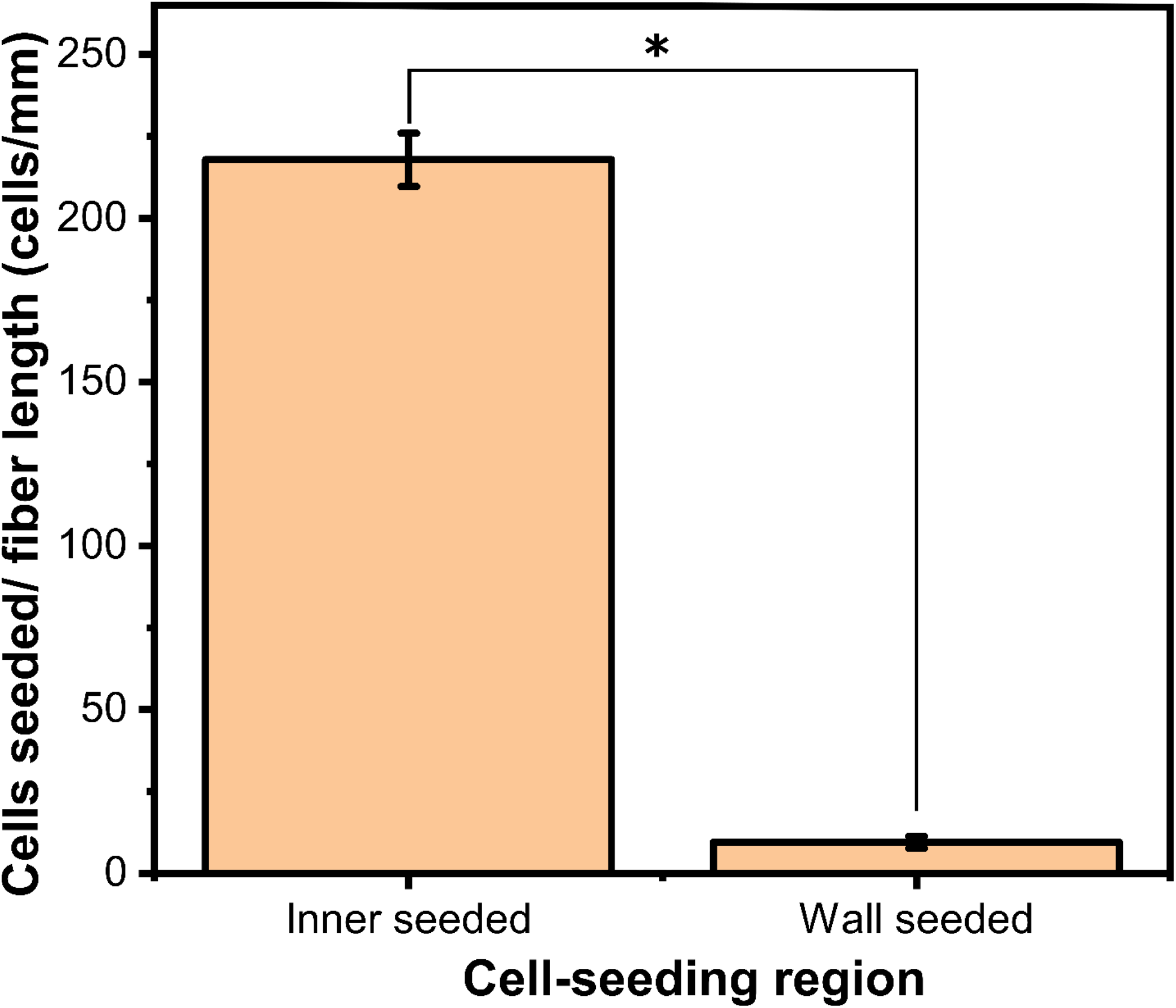
Plot showing the number of cells seeded per mm length of the alginate hollow microfibers in the hollow region compared to their walls and plotted as mean ± sd (n = 6). This indicates that the majority of cells were seeded on the inner surface of the microfibers, and only a few cells were encapsulated in the walls. ANOVA test was performed, and a significant difference was observed (*p < 0.05).

The HBECs on day 0 were found to be uniformly seeded and adhered to the inner surface throughout the length of the alginate hollow microfibers, as shown in Figure 2. Unlike existing surfaces that could be coated with ECM matrix by immersing them in the coating solution, here, the precursor and template solution was exposed to the ECM proteins prior to the formation of the substrate. The use of 0.1% gelatin water as a solvent and ECL mixed with the precursor alginate solution and their prolonged exposure (12 h) to the alginate and PEG created an ECM coating within the walls and on the inner surface of the alginate hollow microfibers upon polymerization. It provided HBECs anchoring points to adhere to the inner surface of the hollow microfibers. After 24 h into the MM, the seeded HBECs exhibited end-feet (processes) initiation, as evident from the day 1 image. The processes initiation was observed to be comparatively slower than the HBECs cultured in the flasks. This could be attributed to the uneven surface of the hollow microfibers, as the HBECs are designed to grow and proliferate on a flat surface of the culture flasks.

Through 6 days in the MM, the HBECs were seen in the process of establishing a monolayer by forming a chain of cells along the length of the microfiber, as shown in **Figure 4**. This chain was observed to initially grow on the edge along the longitudinal direction and eventually progress towards the transverse direction. This behavior could be attributed to the cells migrating towards the edges during the proliferation process. The cell debris was observed to be trapped within the hollow region and can be seen as dark spots. This cell debris could be ascribed to the process of apoptosis and its residue entrapment to the absence of flow within the alginate hollow microfiber. In this study, the HBECs seeded alginate hollow microfibers were vascularized by the MM permeation across the walls of the hollow microfibers. A dynamic system with a solvent flow through the hollow region could wash out the cell debris from the cell-seeded hollow microfibers. Moreover, the shear stress that could be induced on the seeded HBECs due to the flow would also assist in the growth and proliferation of the cells and could be exploited further. The focus of this study was to translate the growth of HBECs from a flat culture flask to a robust inner surface of the alginate hollow microfibers simultaneously during their manufacturing.

**Figure 4.**
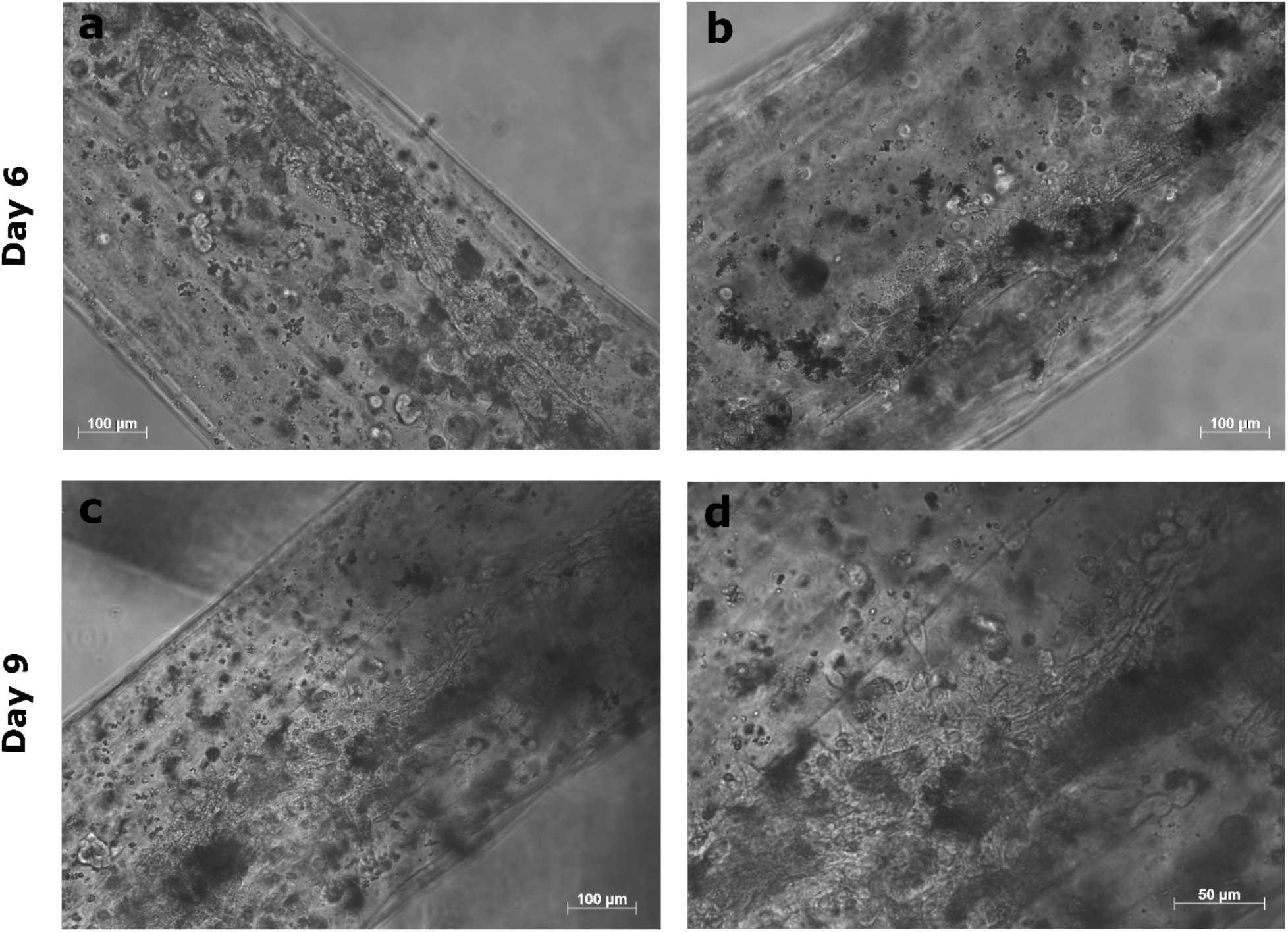
Brightfield images depicting the growth of HBECs on the inner surface of alginate hollow microfibers cultured in the maintenance media. **a** & **b** show the images captured on Day 6 where the chain of HBEC was observed along the inner edge of the hollow microfiber. **c** & **d** show the images captured on Day 9 where the HEBC monolayer was observed in the central region of the hollow microfiber. All scale bars are 100 μm.

The day 9 images of the HBECs seeded alginate hollow microfibers are shown in **Figure 5**. HBECs were observed forming a monolayer on the inner surface of these microfibers, as evident from day 9 images. The HBEC monolayer was observed to be expanded along the width of the microfibers, which was first formed at the edges by day 6 in the MM. A cross-layer connection between two HBEC chains formed on opposite internal surfaces was established. Figure 5 shows the images of the formation of the cross-layer connection between two HEBC chains. The images were captured at different z levels to accommodate the two chains of HBEC formed at the edges of the microfiber. Figure 5 a & b depicts the two layers of the HBEC chains at different z levels, establishing a connection through a cell layer link. The zoomed-in images included in Figure 5 c & d shows one of the intermittent cross-layer connections established in the process of further growing a uniform cell monolayer. This exploratory study demonstrated a unique method to seed and further grow an HBEC monolayer on the inner surface of a hollow alginate microfiber. Future research aims to manufacture HBECs seeded alginate hollow microfibers incorporated with human pericytes and astrocytes encapsulated in the walls to replicate the complete structure of the BBB. Additionally, a platform to mount these cells cladded hollow microfibers will be designed to initiate flow through them, which could subsequently be used to study the perfusion of different xenobiotic compounds across the barrier of the in vitro BBB model. This approach can be extrapolated in seeding several other cell types of microvasculature by choosing a compatible ECM matrix to mimic a range of microvasculature systems in vitro.

**Figure 5.**
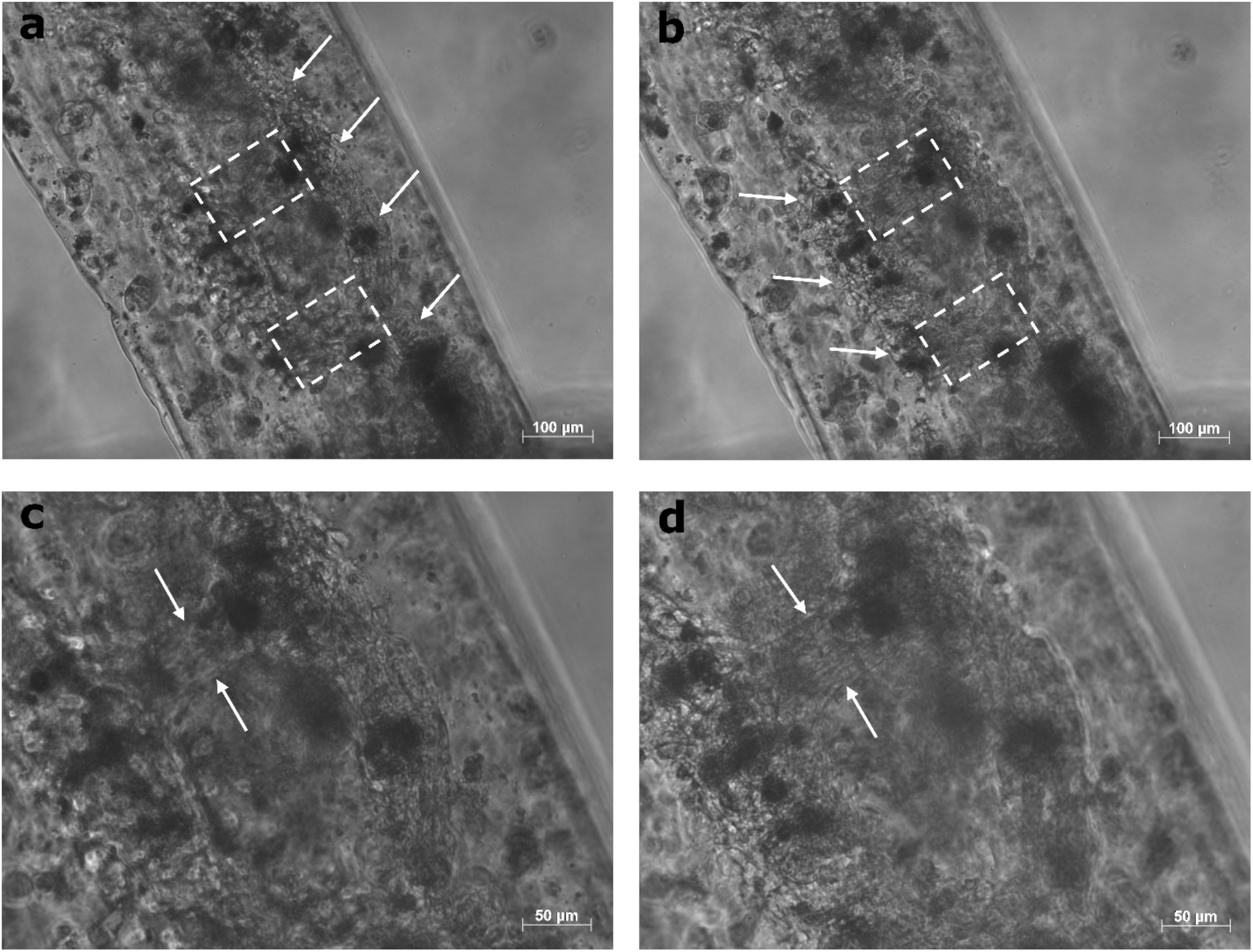
Microscopic brightfield images captured on Day 9 elucidating the progressive formation of HBEC monolayer on the inner surface of alginate hollow microfibers cultured in maintenance media. **a** & **b** show the images of a specific region at different focal length to capture the two chains of HBEC formed at either edge. The white arrows point towards the cell chain in focus. The scale bar is 100 *μ*m. **c** & **d** show the micrographs of HBEC cross-layer connecting the two cell chains. The white arrows point at the cross-layer connection. The scale bar is 50 μm.

## Conclusions

Biomimicking the blood-brain barrier (BBB) in vitro by explicitly replicating its hollow tubular native structure is currently a major challenge. Moreover, establishing a human brain endothelial cell (HBECs) layer on the inner surface of this scaffold adds more to its complexity. This study demonstrated a microfluidic approach to obtain an HBECs monolayer on the inner surface of alginate hollow microfibers by invasively seeding cells simultaneously during the manufacturing process. Results show that the seeded HBECs progressively formed a monolayer on the inner surface of the alginate hollow microfibers by day 9 stored in the maintenance media. A higher cell seeding density of 217 cells/mm length of the microfibers was obtained, which facilitated initial cell growth. The cells were seeded with around 96 % accuracy on the inner surface of the alginate hollow microfibers. The use of gelatin and ECL as an extracellular matrix that coated the surfaces of alginate hollow microfibers during their production facilitated the cell adherence and growth of the seeded HBECs. The demonstrated technique is not confined to only seeding HBECs but could also be extrapolated to seeding different cell types on the inner layer of the alginate hollow microfibers to mimic different microvascular systems in vitro.

## Acknowledgments

This work was partially supported by the National Science Foundation Award 2014346.

